# Liraglutide improves hepatic steatosis and metabolic dysfunctions in a 3-week dietary mouse model of non-alcoholic steatohepatitis

**DOI:** 10.1101/640854

**Authors:** Thibaut Duparc, François Briand, Charlotte Trenteseaux, Jules Mérian, Guillaume Combes, Souad Najib, Thierry Sulpice, Laurent O. Martinez

**Author notes:** Corresponding author: L. Martinez, INSERM UMR1048, Bât. L3, Hôpital Rangueil, BP 84225, 31432 Toulouse cedex 04, France.

## Abstract

**Aim:** Non-alcoholic steatohepatitis (NASH) is an emerging health problem worldwide. However, efficacious pharmacological treatment for NASH is lacking. A major issue for preclinical evaluation of potential therapeutics for NASH is the limited number of appropriate animal models, i.e., models that do not require long-term dietary intervention and adequately mimic disease progression in humans. The present study aimed to evaluate a 3-week dietary mouse model of NASH and to validate it by studying the effects of liraglutide, a compound in advanced clinical development for NASH.

**Methods:** C57BL6/J mice were fed a diet high in fat (60%), cholesterol (1.25%) and cholic acid (0.5%) along with 2% hydroxypropyl-β-cyclodextrin in drinking water (HFCC-CDX diet). Histological and biological parameters were measured at 1 and 3 weeks. Following 1-week diet induction, liraglutide was administrated daily for 2 weeks, and then NASH-associated phenotypic aspects were evaluated in comparison with control mice.

**Results:** Prior to treatment with liraglutide, mice fed the HFCC-CDX diet for 1 week developed liver steatosis and had increased levels of oxidative-stress markers and hepatic and systemic inflammation. For mice not treated with liraglutide, these aspects were even more pronounced after 3 weeks of the dietary period, with additional liver insulin resistance and fibrosis. Liraglutide treatment corrected the diet-induced alterations in glucose metabolism and significantly reduced hepatic steatosis and inflammation.

**Conclusion:** This study provides a novel 3-week dietary model of mice that rapidly develop NASH features, and this model will be suitable for evaluating the therapeutic efficacy of compounds in preclinical drug development for NASH.

## 1 INTRODUCTION

Non-alcoholic fatty liver disease (NAFLD) is a chronic disease that has become pandemic, as it affects between 20% and 30% of the general adult population of most Westernized countries.

Although most cases of NAFLD follow a benign course, approximatively 20% evolve into an aggressive form termed non-alcoholic steatohepatitis (NASH), in which inflammation in the liver can cause hepatocyte damage with or without fibrosis [1]. NASH typically entails high levels of hepatic lipid peroxidation, oxidative stress, apoptosis and production of pro-inflammatory and pro-fibrotic cytokines that induce necroinflammation and ultimately fibrosis [2, 3]. Progressive evolution of hepatic inflammation and fibrosis can lead to cirrhosis, liver failure or hepatocellular carcinoma, with end-stage liver disease due to NASH being anticipated as the leading indication for liver transplantation in subsequent years [4, 5]. Generally, pathogenic drivers and rates of progression of NAFLD are not identical among all patients, and thus NASH should not be thought of as an orderly progression of stages. Although certain advances have led to a better understanding of the pathogenesis of NAFLD and to the identification of therapeutic targets and drugs, no therapy for NASH exists other than lifestyle modifications (to ensure weight loss) and bariatric surgery for morbidly obese patients [6, 7]. Although drug development for NASH has accelerated in recent years, the choice of animal models remains one of the major issues for the assessment of drug candidates at the initial steps of preclinical development. Indeed, an appropriate animal model should recreate as closely as possible the pathological patterns, metabolic and transcriptomic features, and histological alterations found in human NASH. Several preclinical mouse and rat models for NASH are currently available, with various approaches based on genetic modification, e.g., *ob*^−^/*ob*^−^ or *db*^−^/*db*^−^ mice with leptin and leptin receptor deficiency, respectively, diet induction, e.g., high-fat/cholesterol/fructose diet or methionine/choline– deficient diet, or chemical induction, e.g., thioacetamide or carbon tetrachloride [2, 8, 9]. Each of these models can be used separately or in combination. Although these models are mandatory tools for assessing the efficacy of candidate compounds before proceeding to clinical trials, the majority do not entirely replicate human NASH features, i.e., they may lack fibrosis or lack insulin resistance in diet-induced models[10]. Another important issue that may hinder preclinical assessment of NASH drug candidates is the large amount of resources and time needed to bring a model to a standard suitable for experimental studies, as most of the existing models require several weeks, ranging from 6 to 40 weeks, to create a NASH phenotype [2]. Hence, there is need for animal models that rapidly develop NASH features and recapitulate the human pathology as accurately as possible, which would permit an initial, economical, and rapid preclinical evaluation of drug candidates.

To address the issues associated with current NASH models, we developed, characterized, and evaluated the utility of a rapid-onset and inexpensive murine model of NASH based on dietary induction. Toward this goal, we customized a high-fat diet (60% of calories) containing high levels of cholesterol (1.25%) and cholic acid (0.5%), components that have been used separately or in combination to promote hepatic steatosis and lipotoxicity [11–13]. To promote cholesterol uptake by the liver, we also supplemented the drinking water with 2% hydroxypropyl-β-cyclodextrin (CDX), a cyclic oligosaccharide that has high affinity for sterols [14]. Beginning at age 8 weeks, C57BL/6J mice were fed this high-fat, high-cholesterol, cholate and CDX (HFCC-CDX) diet for 1 or 3 weeks, after which parameters related to hepatic steatosis, liver inflammation, liver fibrosis, and glucose metabolism were measured. In parallel, another group of experimental mice were fed the HFCC-CDX diet for 3 weeks, but after 1 week they were also treated daily with liraglutide, a GLP-1 analog currently in advance clinical development for NASH [15]. These groups were then compared with respect to NASH phenotype.

## 2 MATERIALS AND METHODS

### 2.1 Animals, diets and drugs

All animal procedures were performed in accordance with the guidelines of the Committee on Animals of the Midi-Pyrénées Ethics Committee on Animal Experimentation and with the French Ministry of Agriculture license. Wild-type C57BL/6J male mice were caged in animal rooms with 12-h light/12-h dark cycle and *ad libitum* access to water and diet. At the initiation of the dietary intervention, all animals were 8 weeks old and were fed a chow diet (V1534-000, Ssniff, Germany). During the experimental period, mice were provided with either the chow diet for 3 weeks (CD control group) or a custom diet high in fat (60 kcal%), cholesterol (1.25%) and cholic acid (0.5%) (D11061901, Research Diet, Denmark) as well as 2% CDX (Fisher Scientific, France) in drinking water (HFCC-CDX group).

For the last 2 weeks of the 3-week period, groups of mice fed the HFCC-CDX diet received a daily intraperitoneal injection of either liraglutide (Victoza^®^, Novo Nordisk, Denmark) at 100 μg/kg body weight or vehicle (phosphate-buffered saline).

### 2.2 Liver histology

A sample of the main liver lobe was fixed with paraformaldehyde, embedded in paraffin, and sliced into 5-μm sections, then deparaffinized, rehydrated, and stained with Sirius Red or H&E to assess histopathology. For Sirius Red staining, sections were incubated for 10 min in 1% Sirius Red (Sigma, France) dissolved in saturated picric acid and then rinsed with distilled water. For H&E staining, sections were incubated for 30 min in Mayer hematoxylin solution, rinsed with distilled water for 5 min, and then incubated in saturated lithium carbonate solution for 15 s, rinsed again with distilled water for 3 min, and finally placed in 0.5% alcoholic eosin solution for 30 s. Staining with ORO was performed on paraformaldehyde-fixed liver fractions and mounted in Cellpath^™^ OCT Embedding Matrix (Fisher Scientific, France). Samples were sliced in 7-μm thick sections and stained with 0.5% ORO solution in isopropanol for 15 min. The slides were transferred to a 60% isopropanol solution for 1 min, washed with distilled water, and processed for hematoxylin counter staining. For all staining procedures, sections were dehydrated for 15 min with absolute ethanol and incubated with Histoclear^®^ clearing agent (Euromedex, France) before mounting with Distyrene Plasticizer Xylene (DPX) and coverslipping. After staining, slides were scanned with a NanoZoomer 2.0 RS (Hamamatsu, Japan) and then used for blinded histopathological NAFLD activity scoring based on a scoring system adapted from Kleiner *et al*. [16]. Hepatocellular steatosis, liver inflammation, and lobular fibrosis were qualitatively assessed and ranked with a score, and a total NAFLD activity score was then calculated for each animal by summing the scores for steatosis, inflammation, and lobular fibrosis.

### 2.3 Liver lipid content

Total lipids were measured in liver tissue after extraction with chloroform:methanol (2:1) according to Folch *et* al.[17]. Briefly, 100 mg liver tissue was homogenized in 900 μl phosphate buffer pH 7.4 until complete tissue lysis. Lipids were extracted by mixing 125 μl of each lysate with 1 ml chloroform:methanol (2:1). After centrifugation, the chloroform phase was evaporated under nitrogen, and the dried residue was solubilized in 200 μl isopropanol. Triglycerides and cholesterol were measured using commercial kits based on the CHOD-PAP and GPO-PAP detection methods (Biolabo SA, Maizy, France). Results are expressed as micrograms lipid per milligram liver.

### 2.4 Measuring liver ROS and TBARS

ROS in liver were assayed according to the protocol of Szabados *et al*. [18]. Briefly, 30 mg of liver tissue was homogenized in a cold buffer (150 mM KCl, 20 mM Tris-base, 0.5 mM EGTA, 1 mM MgCl_2_, 5 mM glucose and 0.5 mM octanoic acid, pH 7.4). After complete tissue lysis and centrifugation at 12,000 × *g*, the ROS-reactive fluorescent probe H_2_-DCFDA (Molecular Probes) was added to each supernatant to yield 4 μM final concentration of probe, and samples were incubated at 37°C for 30 min with slight agitation. Then, 125 μl of 70% ethanol and 125 μl of 0.1 N HCl were added to stop the reaction. Samples were centrifuged at 3000 × *g* for 15 min at 4°C, and supernatants were collected in new tubes and neutralized by addition of 175 μl of 1 M NaHCO_3_ solution. Each supernatant (200 μl) was added into a black 96-well plate (NUNC), and fluorescence was measured (485 nm excitation, 535 nm emission). Relative fluorescence units were normalized with protein content of each sample, and results are expressed as the percentage of fluorescence versus control.

To assess lipid peroxidation, TBARS, which are byproducts of lipid peroxidation, 200 μl of 0.6% TBA and 10 μl of 1% H_3_PO_4_ solutions were added to 100 μl of liver homogenate. Samples were incubated at 95°C for 1 h, followed by the addition of 200 μl of butanol, mixing, and centrifugation at 12,000 × *g* to extract the colored phase. Fluorescence was measured (515 nm excitation, 548 nm emission), and the TBARS concentration was calculated in comparison with a standard curve. Results were normalized with the protein content of each sample and are expressed as nmol/mg protein.

### 2.5 RNA extraction and real-time quantitative PCR analysis

Total RNA was prepared from liver tissue using QIAzol lysis reagent (QIAGEN, Germantown, USA). Extracted RNAs were suspended in DNase/RNase-free water, and the concentration of each sample was calculated after measuring absorbance at 260 nm using a NanoDrop^™^ spectrophotometer (Thermo Scientific^™^, Waltham, USA). Reverse transcription of mRNA (1 μg) was performed using MMLV Reverse Transcriptase (ThermoFisher, Waltham, USA) in the presence of a random primer (oligo(dT), 2 μl, 500 μg/ml; Promega, Madison, WI, USA) and 0.5 μl RNaseOUT^™^ (ThermoFisher). After 8 min of incubation at 75°C to inactivate DNase, 20 U M-MLV Reverse Transcriptase was added with subsequent incubation at 22°C for 10 min and then at 37°C for 1 h. The reverse transcription reaction was terminated by heating at 95°C for 5 min and then chilling and storing at −80°C.

Real-time quantitative PCR was performed using the SsoFast^™^ EvaGreen^®^ Supermix (Bio-Rad). Expression of each gene was studied and compared with that of the housekeeping gene encoding RPS29 (ribosomal protein 29). PCR was carried out in a 96-well format with 20-μl reactions containing SsoFast^™^ EvaGreen^®^ Supermix (10 μl), cDNA (2 μl), gene-specific primers (0.5 μl) and DNase/RNase-free water (7 μl). Reactions entailed a standard two-step cycling program: 95°C for 3 s and 60°C for 30 s, for 40 cycles. mRNA levels were calculated relative to the average value obtained for the housekeeping gene and further normalized to the relative expression of the respective controls (*i.e*., wild-type mice fed the CD diet).

### 2.6 Analysis of plasma samples

Commercial colorimetric kits (Biolabo SA) based on the CHOD-PAP and GPO-PAP detection methods, coupling enzymatic reaction, and spectrophotometric detection of reaction end-products were used for determination of plasma triglycerides and cholesterol levels. Plasma ALT and AST were determined using a COBAS-MIRA+ biochemical analyzer (Anexplo facility, Toulouse, France).

Levels of inflammatory cytokines (IL6, IL10, TNF-α and MCP-1) were assayed in mouse plasma samples collected by cardiac puncture upon sacrifice; a Milliplex mouse cytokine magnetic kit (Millipore, France) was used according to the manufacturer’s instructions.

### 2.7 Oral glucose tolerance test

After an overnight fast, mice were given an oral gavage glucose load (3 mg • g^−1^ body weight). Blood glucose in samples obtained from the tail vein was assayed with a portable glucometer 30 min before oral glucose loading and at 0, 15, 30, 45, 60, 90 and 120 min thereafter. Insulin concentration in plasma (5 μl sample) was determined 30 min before and 15 min after glucose loading using an ELISA kit (Mercodia, Uppsala, Sweden).

### 2.8 Analysis of insulin signaling

To analyze the liver insulin signaling pathway, mice received insulin at 1 mU • g^−1^ body weight (Actrapid^®^; Novo Nordisk) under anesthesia (ketamine, 100 mg • kg^−1^, Rompun^®^, Bayer, France; and xylazine, 15 mg • kg^−1^, Imalgene 1000^®^, Merial, France) into the portal vein. At 3 min post-injection, mice were euthanized and the liver rapidly dissected. Each individual mouse liver was homogenized using the Precellys lysing kit in RIPA buffer containing a protease inhibitor cocktail (leupeptin, aprotinin). Protein concentration in thawed samples (stored at –80°C) was measured using the Bradford method (Bio-Rad).

The percentage of phospho-Ser-473-Akt to total Akt was determined by capillary western blotting using the ProteinSimple Wes System with 12- to 230-kDa Wes Separation Module capillary cartridges (ProteinSimple, Santa Clara, CA, USA). A mouse monoclonal antibody specific for Akt (#2920S, Cell Signaling) and rabbit polyclonal antibody specific for phosphorylated Akt (pAkt, Ser473, #9271T, Cell Signaling) were used at a dilution of 1:100. Anti-mouse and anti-rabbit detection modules for Wes (ProteinSimple) kits included Luminol-S, peroxide, antibody diluent, streptavidin-coupled horseradish peroxidase, and anti-mouse and anti-rabbit secondary antibodies. Sample proteins (5 ng per each condition) were allowed to separate via the capillary technology and were analyzed based on chemiluminescence, which was transformed into digital images depicting bands as observed in western blots. The abundance of each of total Akt and pAkt was determined using Compass software (ProteinSimple). The normalized data are expressed as the percentage of pAkt to total Akt.

### 2.9 Statistical analysis

Data are expressed as the mean ± SEM. Differences between groups were assessed using the Student’s t-test or one-way analysis of variance followed by Bonferroni *post-hoc* test when necessary. The data were analyzed using Prism V.5.01 for Windows (GraphPad Software, San Diego, CA, USA). The level of significance was defined as p < 0.05.

## 3 RESULTS

### 3.1 HFCC-CDX feeding for 1 week causes hepatic steatosis and inflammation that is associated with systemic inflammation

We first evaluated the ability of the HFCC-CDX diet to induce NASH features within 1 week. C57BL6/J male mice (8 weeks old) were divided into two groups and fed either a chow diet (CD) or HFCC-CDX diet for 1 week. The livers of mice fed HFCC-CDX were grossly enlarged (~30% increase in liver weight; Figure 1A) and pale in color (Figure 1B). Liver histological analyses were performed using Oil Red O (ORO), hematoxylin and eosin (H&E) and Sirius red (SR) staining. ORO staining revealed lipid accumulation in the HFCC-CDX group as compared with the CD group (Figure 1B, ORO), and H&E staining revealed that the HFCC-CDX diet initiated infiltration of mononuclear inflammatory cells, which were absent in the CD-fed group (Figure 1B, H&E, white open circle). Fibrosis was detected in neither groups (Figure 1B, SR). Given the apparently healthy liver phenotype of the CD group, an NAFLD activity score was calculated only for the HFCC-CDX group, and the score for each of steatosis and inflammation confirmed the histological observations (Figure 1C).

**Figure 1:**
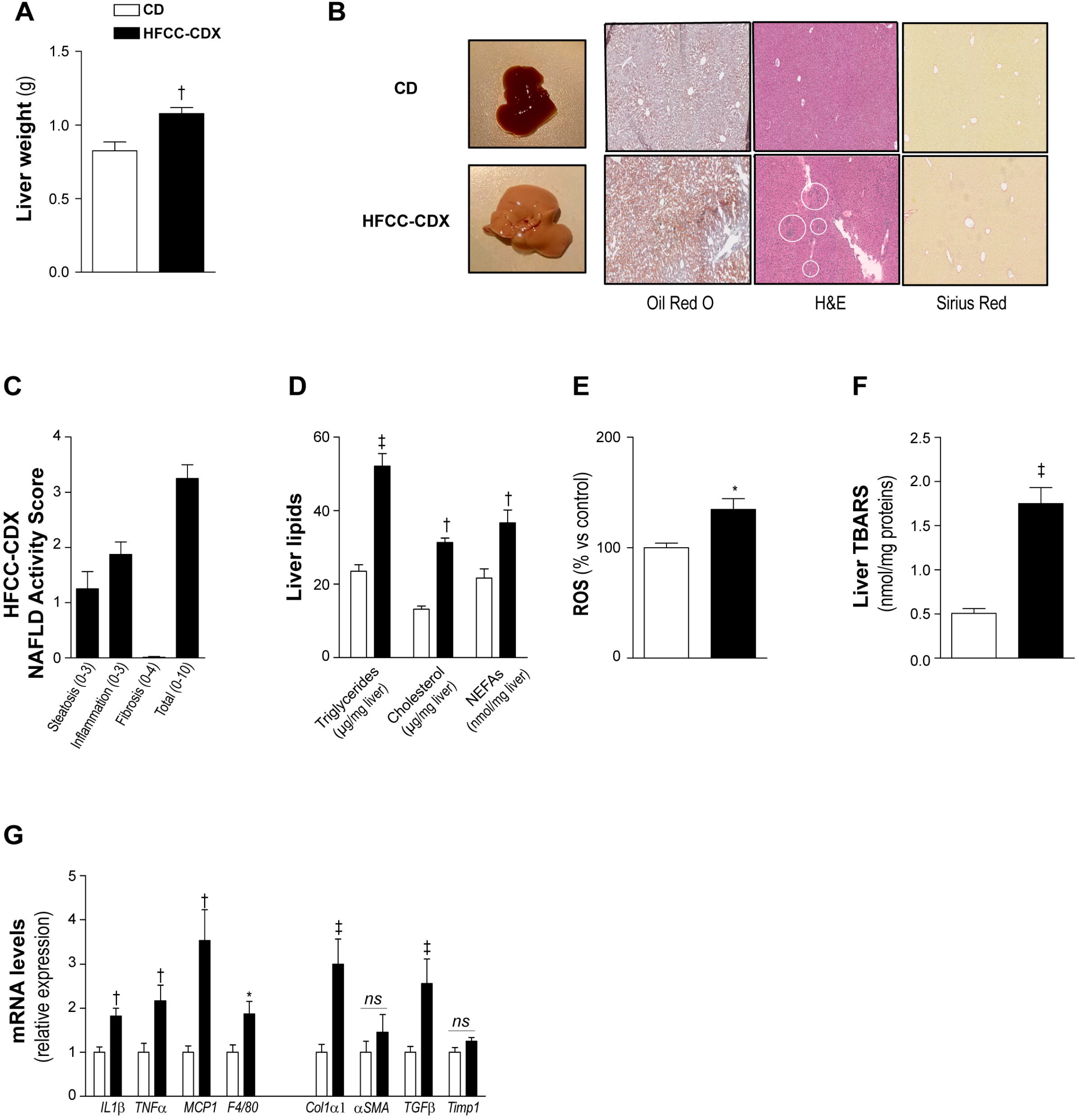
HFCC-CDX diet for 1 week induces liver steatosis, oxidative stress and inflammation. **A)** Liver weight. **B)** Representative photographs of macroscopic liver aspect, Oil Red O, hematoxylin and eosin (H&E), and Sirius Red staining in mice fed a HFCC-CDX or chow diet (CD) for 1 week (magnification 10×). White circles indicate the mononuclear-cell infiltrate (inflammation). **C)** NAFLD activity score for mice fed HFCC-CDX. **D–F)** Assessment of hepatic triglycerides, cholesterol, and non-esterified fatty acids (NEFAs) (D), ROS (E), and TBARS (F). **G)** Expression of pro-inflammatory genes (*IL1β, TNFα, MCP1* and *F4/80*) and pro-fibrotic genes (*Col1α1, αSMA, TGFβ* and *Timp1*) in liver. White bars: chow diet (CD); black bars: HFCC-CDX diet. Results are presented as the mean ± SEM, and the statistical significance of differences was determined with the Student’s t-test. *p < 0.05, ^†^p < 0.01, ^‡^p < 0.001, *ns*: not significant. All data were obtained with 8-week-old mice fed the HFCC-CDX or chow diet (CD) for 1 week, n = 10 per group.

Analysis of hepatic lipid content revealed a ~2-fold increase in each of hepatic triglycerides, cholesterol, and non-esterified fatty acids (NEFAs) as compared with the CD-fed group (Figure 1D), confirming the establishment of hepatic steatosis within 1 week of HFCC-CDX feeding.

Hepatic cholesterol and free fatty acid overload induce oxidative stress that drives lipotoxicity and cellular injury, which are considered critical factors in NASH pathogenesis [19]. We thus analyzed whether the HFCC-CDX diet could induce hepatic oxidative stress and lipid peroxidation by measuring the production of reactive oxygen species (ROS) and thiobarbituric acid response substrates (TBARS), which are byproducts of lipid peroxidation [20]. Both hepatic ROS and TBARS levels were significantly higher in the HFCC-CDX–fed group, with a ~30% and ~300% increase, respectively, compared with the CD group (Figure 1E and 1F). We further analyzed hepatic expression of candidate genes associated with NASH pathology. Confirming the inflammatory response to HFCC-CDX feeding, expression of genes encoding inflammatory cytokines and macrophages chemotactic factors, including *TNFα, MCP-1, IL1β* and *F4/80*, were significantly increased in HFCC-CDX mice compared with CD mice (Figure 1G). Interestingly, the hepatic expression of certain genes related to fibrosis, e.g., *Col1α1* and *TGFβ*, was also significantly increased in the HFCC-CDX group (Figure 1G), suggesting that the fibroproliferative response was also activated.

We then analyzed NASH-related systemic effects. As anticipated, mice fed HFCC-CDX for 1 week had a dyslipidemic profile, with ~200% higher plasma triglycerides and ~60% higher cholesterol levels compared with the CD mice (Figure 2A); also, the plasma levels of the transaminases alanine aminotransferase (ALT) and aspartate aminotransferase (AST) were dramatically increased, reflecting liver injury (Figure 2B). Moreover, the HFCC-CDX mice had significantly higher levels of plasma inflammatory cytokines, including interleukin-6 (IL6, ~600%, p < 0.05) and monocyte chemoattractant protein 1 (MCP-1, ~350%, p < 0.05), whereas a trend toward higher IL10 (~150%) and tumor-necrosis factor α (TNF-α, ~170%) was also observed (Figure 2C). Accordingly, the HFCC-CDX diet dramatically increased the inflammatory index based on differences in inflammatory cytokines levels between mice fed the CD diet or HFCC-CDX diet (Figure 2D).

**Figure 2:**
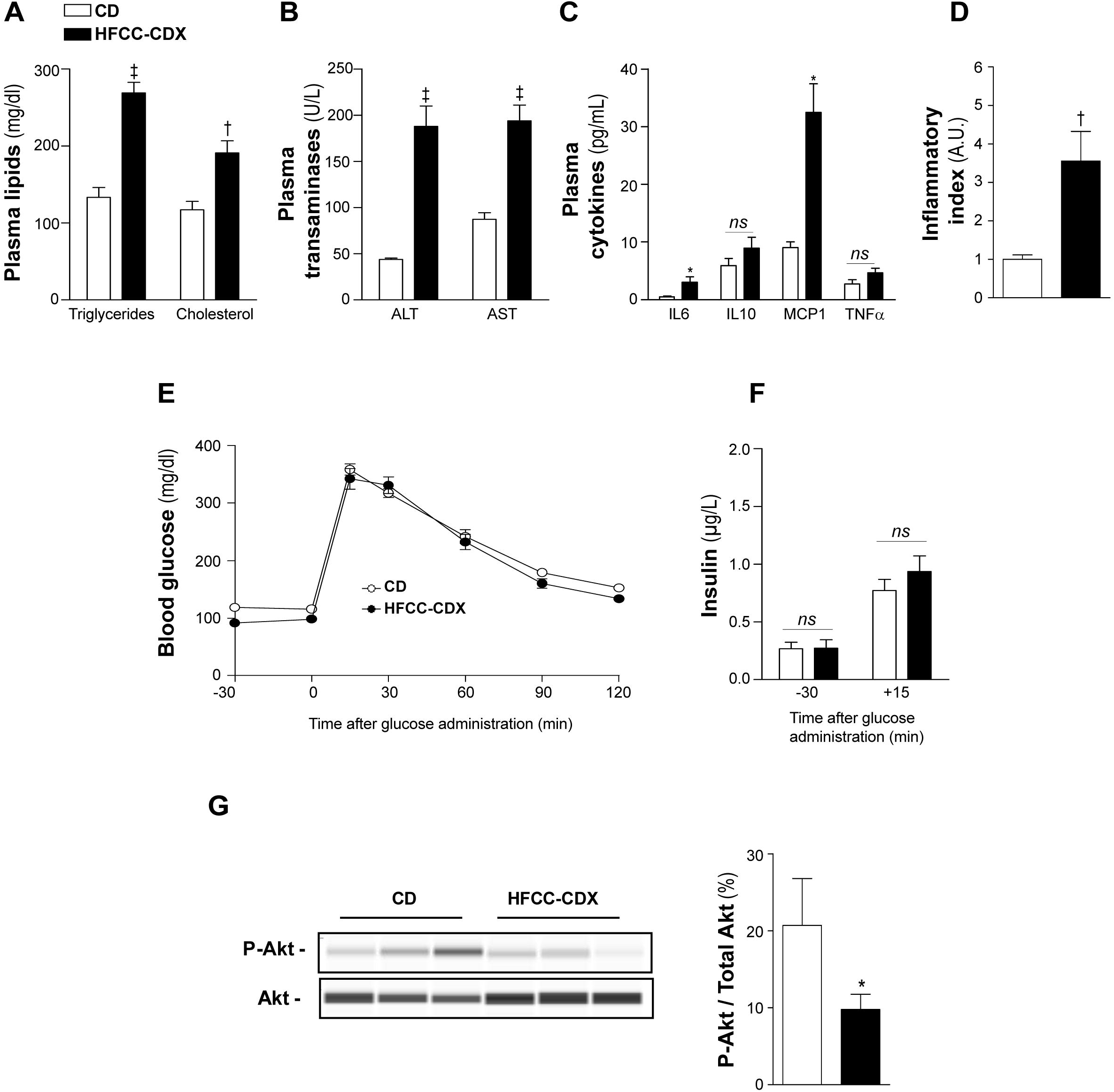
Increase in plasma lipids, systemic inflammation and altered liver insulin sensitivity after 1 week of feeding mice the HFCC-CDX diet. **A)** Plasma triglycerides and cholesterol levels. **B)** Plasma ALT and AST levels. **C)** Plasma concentrations of the inflammatory cytokines IL6, IL10, MCP1 and TNFα. **D)** Index of systemic inflammation based on differences in inflammatory cytokine levels between mice fed the CD diet or HFCC-CDX diet. **E)** Blood glucose evolution after oral glucose loading (oral glucose tolerance test, OGTT) in overnight-fasted mice. **F)** OGTT-associated basal and stimulated insulinemia values in overnight-fasted mice. **G)** Representative western blot of total and phosphorylated Akt levels in the liver of mice after portal insulin injection. Data are expressed as the percentage of phosphorylated Akt to total Akt. White bars: chow diet (CD); black bars: HFCC-CDX diet. Results are presented as the mean ± SEM, and the statistical significance of differences was determined with the Student’s t-test. *p < 0.05, ^†^p < 0.01, ^‡^p < 0.001, *ns*: not significant. All data were obtained using 8-week-old mice fed the HFCC-CDX or chow diet (CD) for 1 week, n = 10 per group.

Next, we found that HFCC-CDX feeding for 1 week did not affect oral glucose tolerance (Figure 2E) and plasma insulin level (Figure 2F). However, direct liver stimulation via injection of insulin into the portal vein caused a significant reduction in phosphorylation of the kinase Akt in the HFCC-CDX group (Figure 2G), reflecting impaired insulin signaling. These results indicated that HFCC-CDX feeding for 1 week established hepatic insulin resistance that had yet to translate into impaired whole-body glucose metabolism. Overall, these data revealed that HFCC-CDX feeding for 1 week is sufficient to induce some important NASH features, including hepatic steatosis and inflammation.

### 3.2 Treatment with liraglutide reverses NASH features and reduces systemic inflammation and insulin resistance induced by the HFCC-CDX diet

To evaluate whether a longer-term dietary intervention could accentuate the NASH features that were observed after 1 week of feeding the HFCC-CDX diet (Figure 1 and 2), a group of mice was fed the diet for two additional weeks (HFCC-CDX group). In parallel, to assess the relevance of this dietary NASH model, we included another group fed the HFCC-CDX diet for 3 weeks to assess whether once-daily treatment with liraglutide for the last 2 weeks could confer protection against NASH [HFCC-CDX-Lira (liraglutide) group]. Liraglutide is a GLP-1 analog licensed for the treatment of type 2 diabetes and is currently in advanced clinical development for NASH [2, 15]. Among all mice used in this experiment, one-third were fed the HFCC-CDX diet and injected with vehicle (phosphate-buffered saline; liraglutide control group) for the last 2 weeks of feeding, and another one-third were fed the HFCC-CDX diet and injected with liraglutide; the other one-third comprised the control mice (no steatosis) that were fed the CD diet.

Total caloric intake was greater for the HFCC-CDX group as compared with the CD group (Figure 3A), but there was no difference in body weight gain between these two groups throughout the feeding period (Figure 3B). As expected, however, liraglutide treatment reduced caloric intake, which was associated with body weight loss compared with non-treated mice (HFCC-CDX group, Figure 3A, B). Compared with HFCC-CDX feeding for 1 week, the NASH phenotype continued to worsen at 3 weeks, as manifested by hepatomegaly (Figure 3C), fatty liver (Figure 3D), and increased content of hepatic triglycerides, cholesterol, and NEFAs (Figure 3E). Treatment with liraglutide significantly reduced liver weight and hepatic triglycerides to a level similar to that of the CD group.

**Figure 3:**
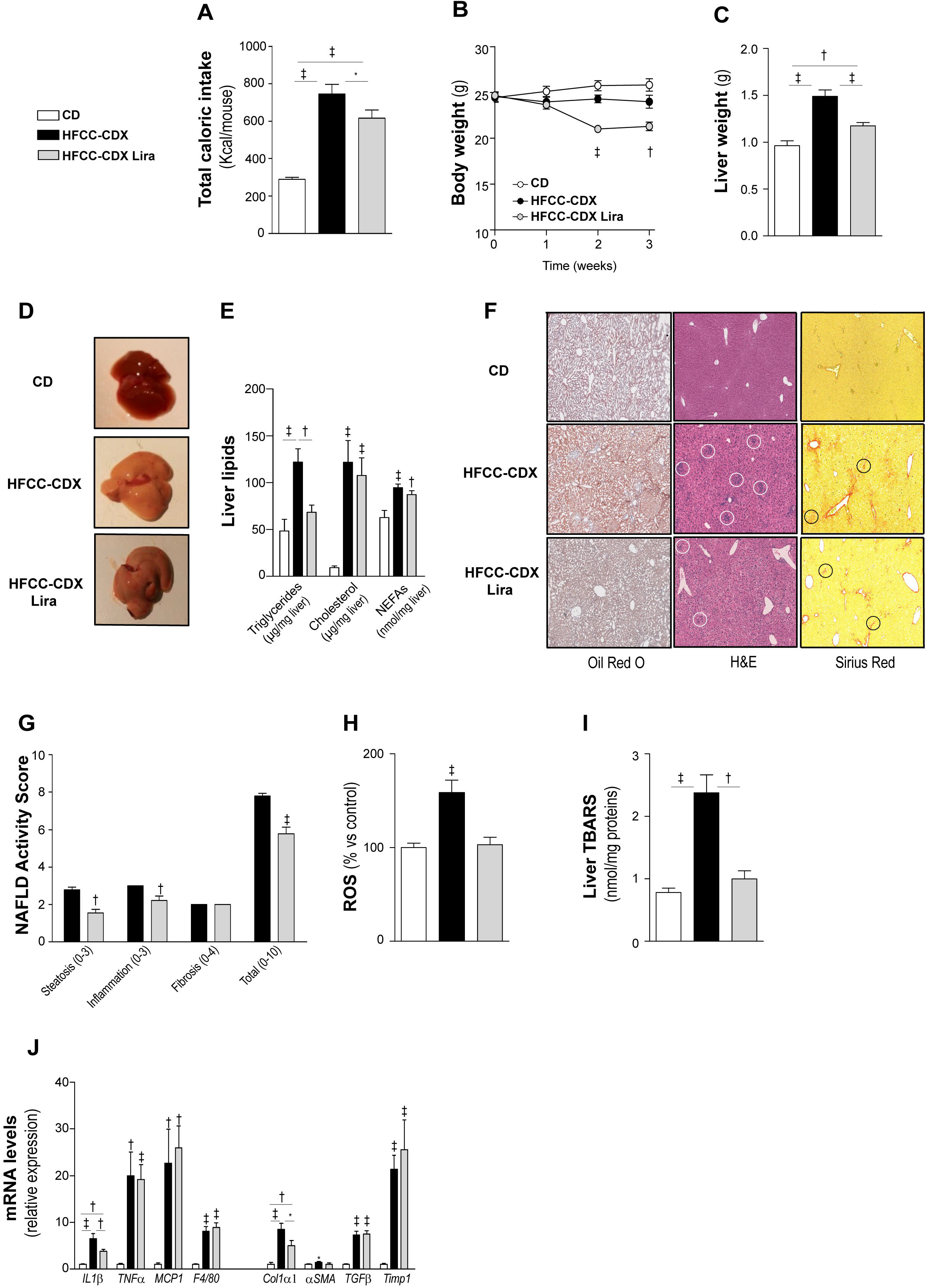
Reversion of liver NAFLD/NASH features with 2 weeks of treatment with liraglutide. **A)** Total caloric intake. **B)** Body weight evolution. **C)** Final liver weight at sacrifice. **D)** Representative photographs of dissected livers at sacrifice. **E)** Content of triglycerides, cholesterol and NEFAs in livers at sacrifice. **F)** Representative images of the histological analysis of livers via staining with Oil Red O, hematoxylin and eosin (H&E), and Sirius Red for mice fed the CD or HFCC-CDX diet and HFCC-CDX–fed mice treated with liraglutide (magnification 10×). White circles indicate the mononuclear cell-infiltrate (inflammation), and black circles indicate collagen deposition (fibrosis). **G)** NAFLD activity score based on the results of staining of liver sections from HFCC-CDX and HFCC-CDX-Lira mice. **H)** Hepatic ROS levels. **I)** Hepatic TBARS measurement. **G)** Liver expression of the pro-inflammatory genes *IL1β, TNFα, MCP1* and *F4/80* and pro-fibrotic genes *Col1α1, αSMA, TGFβ* and *Timp1*. White bars: chow diet; black bars: HFCC-CDX diet; grey bars: HFCC-CDX plus liraglutide. Results are presented as the mean ± SEM, and the statistical significance of differences was determined with the Student’s t-test or with one-way analysis of variance followed by Bonferroni’s *post-hoc* test. *p < 0.05, ^†^p < 0.01, ^‡^p < 0.001; *ns*: not significant. All data were obtained with 8-week-old mice fed a chow diet or HFCC-CDX for 3 weeks. HFCC-CDX–fed mice were treated daily with liraglutide or vehicle for the last 2 weeks of the study, n = 10 per group.

Liver histology was performed (Figure 3F), and the NAFLD activity score was calculated for each of the HFCC-CDX and HFCC-CDX-Lira groups (Figure 3G); the total score comprised the sum of scores for each of hepatocellular steatosis, interstitial inflammation, and lobular fibrosis. For the HFCC-CDX group, steatosis ranged from score-2 limited to Zone-2/3 to a score-3 pan-lobular steatosis. The steatosis was solely micro-vesicular in appearance (Figure 3F, ORO). A maximal score of 3 inflammation was also observed for all animals fed the HFCC-CDX diet. Changes in inflammation were characterized by the presence of a large number of variably sized interstitial foci composed of mononuclear inflammatory cells (Figure 3F, H&E, open white circles). Finally, low score-2 perisinusoidal and peri-portal fibrosis was observed for all animals fed the HFCC-CDX diet (Figure 3F, Sirius Red, open black circles, and Figure 3G). In contrast, mice treated with liraglutide (HFCC-CDX-Lira group) had less extensive and less severe micro-vesicular steatosis and had a reduced number of inflammation-associated mononuclear cells (Figure 3F, G). As with the vehicle-treated mice (HFCC-CDX group), a low score-2 peri-portal/peri-sinusoidal fibrosis was observed for all mice treated with liraglutide. Thus, mice treated with liraglutide had a lower total NAFLD activity score, mainly due to less extensive steatosis and inflammation.

Liver ROS and TBARS levels continued to increase following 3 weeks of HFCC-CDX feeding and liraglutide treatment normalized to values obtained for the CD group (Figure 3H-I), indicating that hepatic oxidative stress and lipid peroxidation induced by HFCC-CDX feeding was resolved by liraglutide treatment.

The ongoing progression of NASH during the 3-week HFCC-CDX diet was confirmed by the dramatic increase in the expression of genes encoding inflammatory cytokines and macrophages chemotactic factors, including *TNFα, MCP-1, IL1β* and *F4/80*, compared with the CD group (Figure 3J), and the differences in expression between the two groups were greater than those observed at 1 week into the study period (Figure 1H). In line with the histological analyses of fibrosis, the expression of genes related to fibrosis (*Col1α1, αSMA, TGFβ* and *Timp1*) were significantly greater in the HFCC-CDX group (Figure 3J). Mice of the HFCC-CDX-Lira group had significantly lower expression of *IL1β, Col1a1* and *αSMA* compared with the HFCC-CDX group.

Concerning NASH-related systemic features, treatment of the HFCC-CDX mice with liraglutide normalized plasma triglyceride levels to those of the CD group, although liraglutide had no effect on the HFCC-CDX–induced hypercholesterolemia (Figure 4A) or on plasma transaminases (Figure 4B). The HFCC-CDX–induced increase in levels of plasma inflammatory cytokines was sustained after 3 weeks of the dietary period, and liraglutide significantly reduced the levels of both IL6 and IL10 (Figure 4C), which translated into a significant reduction of the inflammatory index as compared with the nontreated HFCC-CDX group (Figure 4D). Concerning glucose metabolism, results of the oral glucose tolerance test did not differ between the HFCC-CDX and CD groups (Figure 4E), although the HFCC-CDX diet caused insulin resistance, as shown by the increased basal and glucose-stimulated insulin levels (Figure 4F), and impaired hepatic insulin signaling (Figure 4G). As expected, liraglutide treatment improved glucose tolerance (Figure 4E) and protected from HFCC-CDX–induced insulin resistance (Figure 4F and 4G).

**Figure 4:**
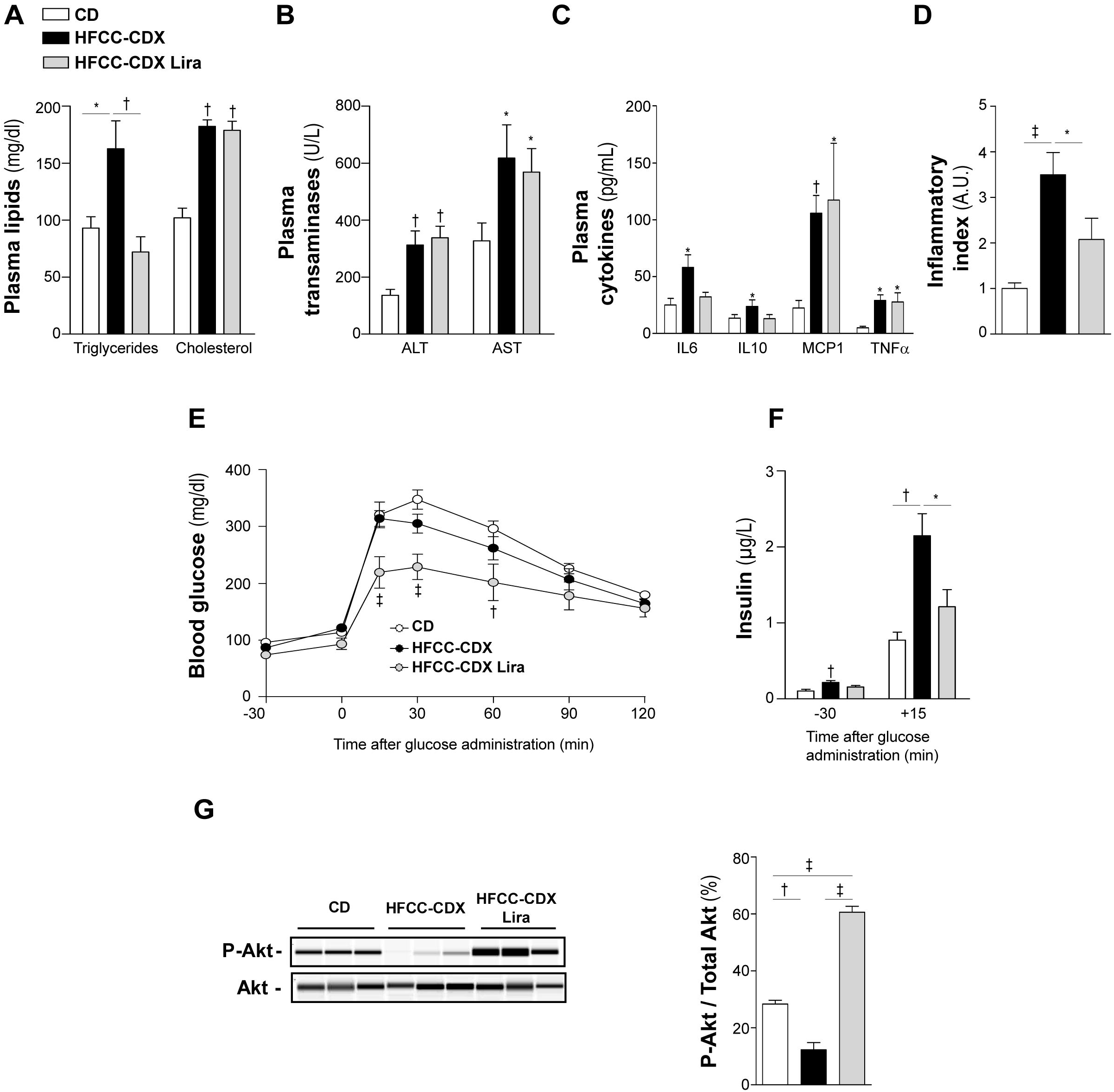
Liraglutide treatment normalizes both plasma lipid levels and systemic inflammation and improves insulin sensitivity. **A)** Plasma triglycerides and cholesterol levels. **B)** Plasma ALT and AST levels. **C)** Plasma concentrations of the inflammatory cytokines IL6, IL10, MCP1 and TNFα. **D)** Index of systemic inflammation. **E)** Blood glucose evolution after oral glucose loading (OGTT) in overnight-fasted mice. **F)** OGTT-associated basal and stimulated insulinemia values in overnight-fasted mice. **G)** Representative western blot of total and phosphorylated Akt levels in the liver of mice after portal insulin injection. Data are expressed as the percentage of phosphorylated Akt to total Akt. White bars: chow diet; black bars: HFCC-CDX diet; grey bars: HFCC-CDX liraglutide. Results are presented as the mean ± SEM, and the statistical significance of differences was determined with the Student’s t-test or with one-way analysis of variance followed by Bonferroni’s *post-hoc* test. *p < 0.05, ^†^p < 0.01, ^‡^p < 0.001; *ns*: not significant. All data were obtained with 8-week-old mice fed the chow diet (CD) or HFCC-CDX diet for 3 weeks. HFCC-CDX–fed mice were treated daily with liraglutide (HFCC-CDX-Lira) or vehicle (HFCC-CDX) for the last 2 weeks of the study, n = 10 per group.

## 4 DISCUSSION

For purposes of preclinical assessment, we characterized and validated a novel dietary mouse model that recapitulates the major hallmarks of human NASH within 3 weeks of the diet. In the proposed model, the C57BL/6 genetic background was chosen because it is more susceptible to liver injury than other inbred strains [21–23]. The HFCC-CDX diet used in this model was formulated to be high in fat (60 kcal%), cholesterol (1.25%) and cholic acid (0.5%) and was supplemented with 2% CDX in drinking water, with the objective of potentiating NASH development *via* the ability of cholic acid and CDX to stimulate the intestinal absorption of cholesterol and lipids and to increase hepatic cholesterol content and inflammation[11, 12, 14]. The NASH features reproduced in this model include liver steatosis (characterized by excessive accumulation of triglycerides, cholesterol and NEFAs within the first week of the diet), a marked rise in the level of the liver-impairment marker ALT, oxidative stress, and hepatic inflammation and progression to mild fibrosis within 3 weeks. This model also exhibited transcriptomic and metabolic alterations associated with the human disease, including insulin resistance, dyslipidemia, and systemic inflammation.

For a short period of dietary intervention (e.g., 3 weeks in our study), none of the current diet-induced mouse models of NAFLD/NASH can recapitulate the human pathogenesis as well our HFCC-CDX model. For instance, feeding C57BL/6J mice a high fat (60% kcal) diet containing cholesterol (1.25%) and cholate (0.5%) for 12 weeks was found to induce steatosis, inflammation, and fibrosis associated with dyslipidemia, lipid peroxidation, oxidative stress, and insulin resistance [24], whereas we found that a similar NASH phenotype was induced in just 3 weeks of feeding with the HFCC-CDX diet. Likewise, Western-type diets of varying composition with respect to fat (21–45 kcal%), cholesterol (0.1–2%), and sugars (e.g., 20% fructose) closely mimic the pathogenesis of human NASH, i.e., insulin resistance, inflammation and liver fibrosis; regardless of the composition, however, the NASH phenotype manifested only after a relatively long-term diet duration of 8 to 30 weeks [2, 9]. C57BL/6J mice fed a methionine- and choline-deficient diet develop a NASH phenotype in the shortest time compared with all previous NASH models, with hepatic steatosis by 2–4 weeks and progression to inflammation and fibrosis shortly thereafter, although animals lose weight and do not develop insulin resistance [25, 26]. In contrast, the present HFCC-CDX dietary model more closely recapitulates the NASH-associated metabolic dysfunctions of humans, including insulin resistance. One limitation of the HFCC-CDX dietary model for NASH compared to the disease development in humans is that mice do not become obese.

We also used our 3-week HFCC-CDX dietary mouse model to test the therapeutic efficacy of liraglutide. Indeed, besides the attributes of improving glucose metabolism, reducing food intake, and promoting weight loss, certain other systemic and hepatic effects of liraglutide have been documented in preclinical and clinical studies, and hence liraglutide has been the subject of extensive evaluation as a therapeutic candidate for NAFLD [27, 28]. Our present study revealed that once-daily injection of liraglutide for 2 weeks decreased caloric intake and body weight, improved glucose tolerance, and protected from HFCC-CDX–induced insulin resistance, which are well-known effects of liraglutide in humans [27, 28]. In addition, we observed that liraglutide could resolve hypertriglyceridemia and reduce hepatic steatosis, oxidative stress, and systemic and hepatic inflammation but not fibrosis, results that are in general agreement with the effects of liraglutide documented in preclinical studies with mice [29–31] and clinical trials for NASH [32–34]. In NAFLD patients, plasma ALT and AST levels have been found to decrease gradually during liraglutide treatment [32–34]. In our experimental setting, liraglutide could indeed significantly decrease the total NAFLD activity score but could not counter the HFCC-CDX–induced increase in plasma ALT and AST, in agreement with results from another dietary mouse model of NASH [30]. Whether longer-term treatment with liraglutide can reduce transaminase levels in mice — as observed in human studies — will require a different experimental design. In this respect, the objective of the present work was to provide a preclinical model to facilitate rapid screening of therapeutic targets to treat NASH, but future studies must explore whether HFCC-CDX fed over a longer period might promote progression to more severe stages of NASH, such as advanced fibrosis, cirrhosis, and hepatocellular carcinoma. Also, we chose the mouse strain C57BL/6 based on its reasonable cost and susceptibility to diet-induced liver steatosis compared wither other inbred strains [21–23]. Future studies could investigate the effect of the HFCC-CDX diet on isogenic strains selected for their susceptibility to develop diet-induced NASH or genetic mouse models that are predisposed to developing a fatty liver [35–37].

In summary, although drug-development research for NASH is intense and advancing rapidly, here we propose HFCC-CDX as a new dietary model of NASH that reproduces the main features of disease development in humans for the purpose of facilitating the rapid screening of drug candidates and prioritizing the more promising candidates for advanced preclinical assessment and subsequent clinical trials.

## ACKNOWLEDGMENTS

The authors thank the following individuals for excellent technical assistance: the staff of the Rangueil animal facility (CREFRE, Toulouse), Lucie Fontaine (Histology core facility, I2MC, Toulouse), Alexandre Lucas (We-Met *Functional Biochemistry Facility*, I2MC, Toulouse) and Laurent Montbrun (ANEXPLO Phenotyping facility, CREFRE, Toulouse).

